# Sequential delay and probability discounting tasks in mice reveal anchoring effects partially attributable to decision noise

**DOI:** 10.1101/2021.06.08.447620

**Authors:** Gerardo R. Rojas, Lisa S. Curry-Pochy, Cathy S. Chen, Abigail T. Heller, Nicola M. Grissom

**Author notes:** to whom correspondence should be addressed: Nicola Grissom, Department of Psychology, University of Minnesota, 75 East River Rd, Minneapolis, MN 55455.

## Abstract

Delay discounting and probability discounting decision making tasks in rodent models have high translational potential. However, it is unclear whether the discounted value of the large reward option is the main contributor to variability in animals’ choices in either task, which may limit translatability to human discounting data. Male and female mice underwent sessions of delay and probability discounting in sequence to assess how choice behavior adapts over experience with each task. To control for “anchoring” (persistent choices based on the initial delay or probability), mice experienced “Worsening” schedules where the large reward was offered under initially favorable delay or probability conditions that became less favorable during testing, followed by “Improving” schedules where the large reward was offered under initially unfavorable conditions that improved over a session. During delay discounting, both male and female mice showed elimination of anchoring effects over training. In probability discounting, both sexes of mice continued to show some anchoring even after months of training. One possibility is that noisy action selection could contribute to these anchoring effects, rather than persistent fluctuations in value discounting. We fit choice behavior in individual animals using models that included both a value-based discounting parameter and a decision noise parameter that captured variability in choices deviating from value maximization. Changes in anchoring behavior over time were tracked by changes in our decision noise parameter, not the value parameter. Thus, changes in discounting behavior in mice can result from changes in exploration of the environment rather than changes in reward valuation.

## 1. Introduction

Delay discounting tasks measure value assessments against a temporal cost, while probability discounting tasks measure value assessments against risky reward (Green & Myerson, 2004; Odum, 2011b). These tasks have been important tools in assessing dysregulated reward processing in neuropsychiatric disorders such as addiction or neurodevelopmental disorders (Andrade & Petry, 2012; Dalley et al., 2011; Richards et al., 1999; Rung, Peck, et al., 2019). Because of this, translational animal versions of these tasks are of high interest (Mitchell, 2014; St Onge & Floresco, 2009). However, it is unclear if animals use similar discounting strategies to those used by humans (Vanderveldt et al., 2016), for two reasons.

One issue arises from the fact that behavior in each of these tasks alone are thought to reflect choice impulsivity in animals (Acheson et al., 2006), even though these tasks contribute in different ways to assessing an impulsive profile (Strickland & Johnson, 2020). It has recently been shown in humans that multiple distinct discounting tasks are needed to better capture common traits (Białaszek et al., 2019); methods testing both delay and probability in the same animals are therefore of strong interest, but not widely available or used, especially in mice.

A second issue is that discounting tasks are typically modeled in both humans and rodents using economic value functions (e.g., k-values and h-values) which assume the main relevant factor in choices is the current discounted value of the reward (Odum, 2011b). However, recent evidence from the literature on reinforcement learning and decision making strongly implicates choice history and exploration as important variables in how both humans and animals perform value-based decision tasks (Chen, Ebitz, et al., 2020; Cinotti et al., 2019; Daw et al., 2011; Speekenbrink & Konstantinidis, 2015). Importantly, animals often engage in non-reward seeking behaviors that are typically described as exploration (Findling et al., 2019; Gershman, 2019). This decision “noise” is rarely considered as a contributor to choices in discounting tasks despite exploratory events being necessary for animals to learn new task rules (Ebitz et al., 2018; Ebitz et al., 2019). In humans, exploration in other decision making tasks correlates with the degree of delay discounting shown (Sadeghiyeh et al., 2020). We have recently identified exploration as a key driver of sex differences in other decision making tasks (Chen, Ebitz, et al., 2020; Chen, Knep, et al., 2020). Because discounting tasks are often used to compare groups of animals modeling neuropsychiatric risk factors or other individual differences such as sex differences (Grissom & Reyes, 2019; Orsini et al., 2016; Orsini & Setlow, 2017; van den Bos et al., 2013; Weafer & de Wit, 2014), it is imperative to identify methods that allow us to distinguish whether differences in behavior are due to value judgements putatively reflecting impulsivity or if exploration is a strong latent contributor to behavior.

One way to address these issues is to develop a method allowing within-subjects comparison of delay and probability discounting functions. This approach would allow for comparing overall performance within and between groups and permit computational modeling across tasks incorporating a decision noise parameter in addition to a value parameter. In the present study, we describe a sequence of delay and probability discounting tasks in mice achieving these goals. Mice are increasingly used for cognitive task batteries because of their high genetic tractability. Recent advancements in technology for mouse operant testing available through touchscreens have substantially improved the ease of training mice (Horner et al., 2013), enabling us to develop matched versions of probability and delay discounting for mice.

Here, we describe a novel battery of sequential delay and probability discounting schedules in touchscreens tested in male and female wildtype mice. One key factor previously shown to affect choices in these tasks is the order of presentation of delays or probabilities on the large reward (e.g. “improving” or “worsening”) (Koffarnus et al., 2013; Maguire et al., 2014; St Onge et al., 2010). We alternated mice between worsening and improving schedules within each discounting task, and demonstrated that these order effects are substantial in both male and female mice. These order effects are fully eliminated in both sexes with extensive training in delay discounting. However, order effects remain in probability discounting, especially in female mice, supporting persistent sex differences in risk processing but not “impulsivity” *per se* (Grissom & Reyes, 2019). We analyzed reward strategy via both win-stay/lose-shift analyses and model fits for both value and decision noise parameters in order to better understand how mice adapt choice strategies between delay and probability discounting. Win-stay/lose-shift analyses suggest sex differences in probability discounting emerged because females learned to win-stay consistently over the course of probability discounting training. Choice parameters showed mice preferred exploration strategies when rewards were delayed and exploitation strategies when rewards were risky. These results demonstrate that value functions may capture one aspect of impulsivity (i.e. overall reward preference), but cannot account for how choice behavior changes between tasks or across multiple experiences of the same task.

## 2. Materials and methods

### 2.1 Subjects

8 male and 7 female BL6129SF1/J mice (from Jackson Laboratories) took part in both delay discounting and probability discounting. Mice began experiments at approximately 70 days of age. Mice had free access to water and were food restricted with their home chow at 85-90% of their baseline weight. Mice were pre-exposed to the operant reinforcer, vanilla flavored Ensure, in their home cage for one day prior to training. On this day, mice were verified to have consumed a full bottle of Ensure (148 ml). Behavioral testing took place Monday to Friday, and on Fridays, mice had free access to home chow. Animals were housed on a reverse light dark cycle (9am-11pm) and were tested during the dark period. Animals were cared for in accordance with National Institute of Health guidelines and were approved by the University of Minnesota Institutional Animal Care and Use Committee.

### 2.2 Apparatus

16 identical triangular touchscreen operant chambers (Lafayette Instrument Co., Lafayette, IN) were used for training and testing. The touchscreen was housed in the front while the food delivery magazine in the back. Information on individual touches on touchscreens throughout sessions were recorded via ABET-II software. Touchscreens were limited by masks with holes which allowed responding in 5 square holes. Liquid reinforcer (50% diluted Ensure) was pumped via a peristaltic pump (1000 ms or 250 ms duration, corresponding to volumes of approximately 25 μl and 6.25 μl). ABET-II software (Lafayette Instrument Co., Lafayette, IN) was used to program operant schedules and analyze all data from training and testing.

### 2.3 Behavioral procedures

#### Magazine training

Mice received free 7 μl of Ensure every 30 seconds for 30 minutes in operant chambers for 5 days. Mice learned to approach the magazine to obtain Ensure.

#### Center Hole Fixed-Ratio 1

Mice were initially trained to nosepoke the center hole of a 5-hole mask on the touchscreen chamber on a Fixed-Ratio 1 schedule for 10 days, 30 minutes each day. 7 μl of Ensure was delivered immediately following a nosepoke. Only the center hole was illuminated during these sessions. The tray holding the Ensure was illuminated until mice interrupted an infrared beam when their head entered the reward port, and this allowed them to move to the next trial.

#### Chaining Center to Left and Right

For this phase of training, hole 3 (center) illuminated and a nosepoke resulted in 7 μl of Ensure. After a center nosepoke and its reward, on the next trial, holes 2 (left) and 4 (right) illuminated, and mice learned that nosepoking one side resulted in a small amount (7 μl) or a large amount (28 μl) of Ensure. Training lasted 29 days with 30 minute sessions. Responses on holes 1 and 5 were counted as non-reinforced touches.

#### Responding on Sides Only

Mice learned to nosepoke only the left and right holes for Ensure at the same volumes as above for 13 days. The left and right holes were the only holes to illuminate during these sessions. Center hole nosepokes no longer delivered the reinforcer. Sessions ended after 60 reinforced trials or 30 minutes had elapsed.

#### Responding on Chained Sides Followed by ITI

Mice then learned to chain a center hole nosepoke to a left or right hole nosepoke for 13 days. The action sequence of center-to-left or center-to-right led to a large reward (25 μl) or small reward (6.25 μl). This phase of the testing was counterbalanced; half of the mice experienced the large reward on the left and the other half on the right. An inter-time interval (ITI) of 30 seconds followed in order to suppress the reward rate. Animals were limited to no more than two trials in a row selecting one side before being forced to try the other side, to ensure they experienced the small reward side as well as the large reward. Sessions ended after 30 minutes had passed.

#### Improving and Worsening Delay Discounting

To test the influence of anchoring effects, we tested mice on a Worsening schedule that was at the start of the session was initially favorable and became unfavorable as the session proceeded, then reversed the order of delays for the Improving schedule, then reversed again 4 additional times. Mice underwent 10 days of Worsening delay discounting I followed by 8 days of Improving delay discounting I, 9 days of Worsening delay discounting II and 10 days of Improving delay discounting II, and 16 days of Worsening delay discounting III and 16 days of Improving delay discounting III. One side delivered a large but delayed reward (25 μl) or a small and immediate reward (6.25 μl). The side with the large reward was matched to the chained side training. If mice responded for the small reward, an ITI occurred based on the delay for that session. If mice chose the large reward, the center hole blinked for the duration of the delay. On the Worsening schedule, mice experienced 12 trials of increasing delays – 0s, 4s, 12s, 20s, 28s delays – within one session. The Improving schedule was similar but in the reverse orientation – 28s, 20s, 12s, 4s, 0s delays – within one session. Mice could only move on to the next delay if they responded on all 12 trials for that delay and collected the reward.

#### Worsening and Improving Probability Discounting

Animals transitioned directly from the last delay discounting schedule to the first probability discounting schedule because the structure of the task (location of large and small reward, response order) did not change, only the rule governing payout of the large reward. Mice underwent 9 days of Worsening probability discounting I and followed by 9 days of Improving probability discounting I, 8 days of Worsening probability discounting II and 9 days of Improving probability discounting II, and 16 days of Worsening probability discounting III and 21 days of Improving probability discounting III. As in delay discounting, the same side resulted in a large reward or small reward. Both rewards were immediate, but the large reward was delivered probabilistically. On the Worsening schedule, mice experienced 20 trials of decreasing probability of reward delivery – 100%, 75%, 50%, 25%, 12.5% chance of reward – within one session. The Improving schedule was similar but in the reverse orientation – 12.5%, 25%, 50%, 75%, 100% probabilities – within one day. If the trial was not rewarded, a feedback house light would blink indicating a non-rewarded trial. Mice could only move on to the next probability if they responded to all 20 trials for that probability and collected the reward.

### 2.4 Computational modeling

To quantitatively examine how the value of rewards varies as a function of delay, we fit an exponential discounting model (Odum, 2011b), shown as in the equation below:

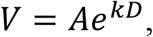

where *V* is the subjective or discounted value of the delayed reward, *A* is the amount or magnitude of the delayed reward, and *D* is the length of delay. *k* is a free parameter that reflects the discounting rate: the larger *k* is, the steeper the discounting of reward value; the smaller *k* is, the slower the discounting of reward value. *k* is determined by the fit of the model to the actual data.

Then, we fit a similar exponential model (Richards et al., 1999) to examine how uncertainty of reward affects the value of reward.

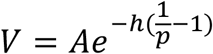

In this model, *p* is the probability of reward and *A* is the magnitude of the reward. The free parameter *h* dictates how steep is the change in the value of reward as a function of the reward probability. Thus, *h* in the probability discounting model and *k* in the above delay discounting model both describe how rapidly the value of a reward is discounted, either by uncertainty or delay of the reward.

For both models, the action selection was performed based on a Softmax probability distribution:

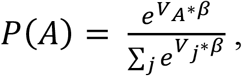

Where V_A_ corresponds to the subjective reward value of action A, and a second free parameter inverse temperature β determines the level of decision noise. When inverse temperature is high, the decision noise is low, which means more exploitation of the action with high subjective value; when inverse temperature is low, the decision noise is large, which means more random exploration regardless of value. The optimized parameters were obtained through minimization of the negative log likelihood of the models.

### 2.5 Statistical analysis

Delay discounting and probability discounting data were analyzed using R Studio using the lme4 package. Linear mixed models were fit to preference proportion data for both delay discounting and probability discounting with fixed effects of sex (males and females) and delay/probability and random effect for subjects (**Figures 2 & 3**). In cases where sex was not significant based on a log-likelihood ratio test, sex was removed from the linear mixed model.

Win-stay scores were calculated by taking the number of times animals stayed on the large risky side after receiving Ensure on the previous trial divided by the total number of times animals received Ensure on the large side. Lose-shift scores were calculated by taking the number of times animals didn’t receive Ensure on the large side and switched to the small certain side divided by the total number of times animals didn’t receive rewards on the large side (regardless if they shifted or not). These scores were then analyzed using linear mixed models with fixed factors of sex and schedule (Improving vs Worsening) and random effect for subjects (**Figure 4**).

Computational modeling results were fit to discounting and inverse temperature parameters (**Figure 5**) and were analyzed with linear-mixed models with task and schedule as the fixed factors and random effects for subjects. For all statistics, an alpha of 0.05 was used.

## 3. Results

Age-matched male and female wildtype mice (n=16, 8 males, 7 females, strain B6129SF1/J) were trained to perform sequential discounting tasks using touchscreen operant chambers (**Figure 1A**). This permitted us to test the extent to which choice preferences and discounting were “anchored” by the initial delay/probability of the large reward. Trials were paced to require 30 seconds minus the length of the delay before the next trial could be initiated to remove the ability to complete all trials more quickly (Pearson et al., 2010) that may contribute to prior demonstrations of greater action impulsivity. Repeated sessions of delay and probability discounting allowed us to study anchoring effects and choice strategy, as well as ask questions about sex differences and individual differences in preferences for delay and probability simultaneously.

**Figure 1.**
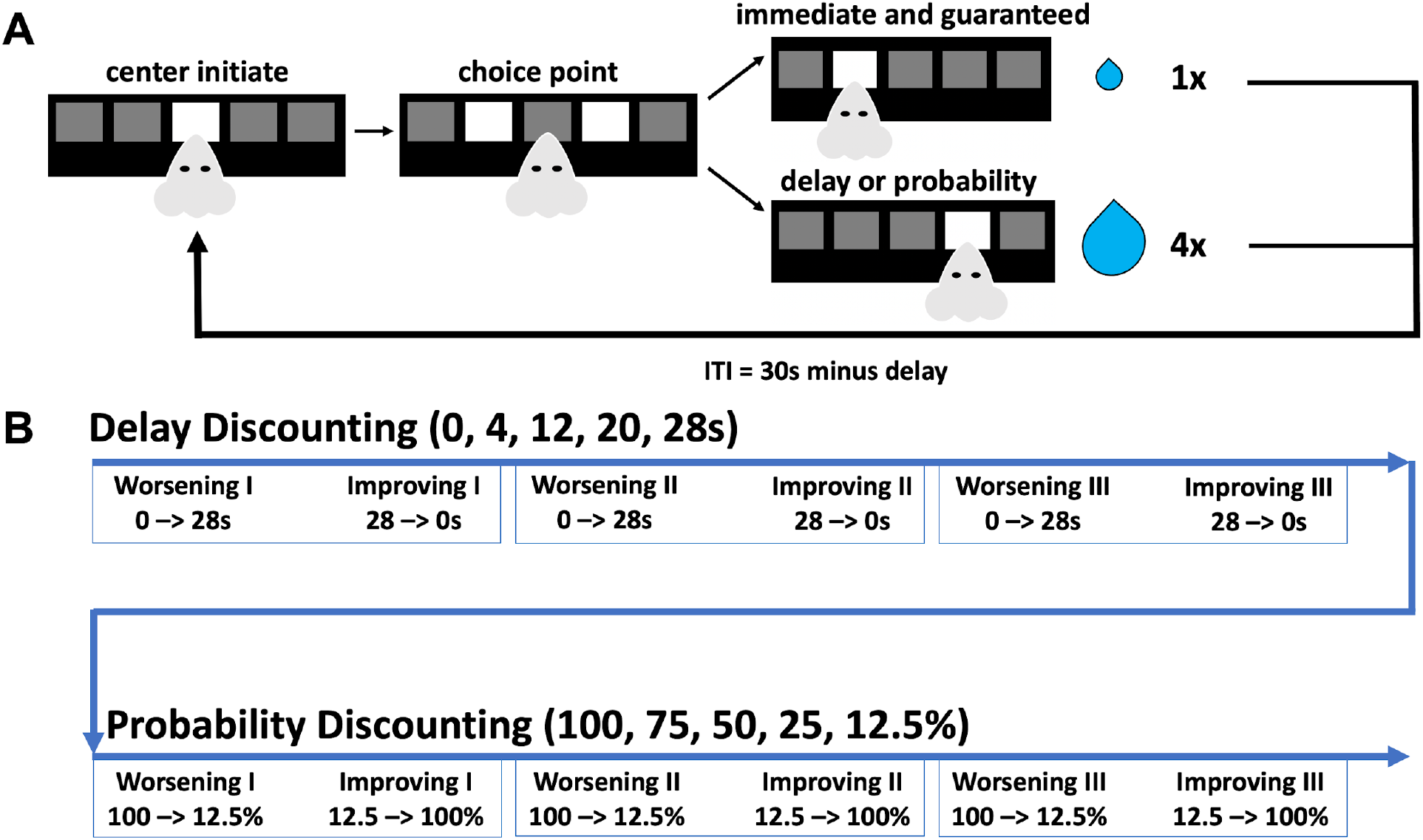
Schematic of delay discounting and probability discounting. **A.** Mice responded for large and small rewards after initiating the trial by poking the center hole. Mice then chose one side for a small immediate reward (6.25 μl) or a large delayed/probabilistic reward (25 μl). Small choices on the delay schedule were followed by an ITI to time match the delay of the large reward. **B.** Mice began discounting tasks on the Worsening schedule (i.e. increasing delay) for 10 days and then were switched to the Improving schedule (i.e. decreasing delay) for 9 days. Schedules continued to switch two more times. Mice then experienced probability discounting on the Worsening schedule (i.e. increasing uncertainty) for 9 days and then were switched to the Improving schedule (i.e. decreasing uncertainty) for 9 days. Schedules again switched two more times.

### 3.1 Delay Discounting

#### Anchoring effects to delayed rewards are reduced with experience, but take longer to do so in females compared to males

Rodent models of delay discounting have previously used latin square design of delays (Mitchell, 2014), but structured delay schedules have not widely been used in mice (e.g. ascending or descending schedules). Because of the novelty of these schedules, we initially put mice through delay discounting in order to test whether sex is an important factor in anchoring effects induced by shifting schedules (i.e. “Worsening” and “Improving”). Research suggests uncertainty in discounting tasks can produce sex-specific effects (Grissom & Reyes, 2019), thus we wanted to test whether sex was important in anchoring responses to Worsening and Improving schedules. Here, we present the data from each round of Worsening and Improving schedules grouped together.

The first time mice experienced both the Worsening and Improving Delay Discounting schedules (Delay Discounting I), their preferences for the large reward across the entire sessions were heavily anchored by the initial delay experienced (**Figure 2A**, main effect of schedule, F_(1, 15.02)_ = 32.44, p < 0.001). Evidence suggested males and females might differently process Worsening and Improving delays (**Figure 2A**, sex x schedule x delay interaction, F_(4, 1076.05)_ = 3.4237, p = 0.0086). Male mice immediately reduced their responding for large rewards on the Improving schedule compared to the Worsening schedule (28s delay, p = 0.0142). Male mice continued to reduce their responding except for when large rewards were no longer delayed (20s delay, p = 0.0182; 12s delay, p = 0.0012; 4s delay, p = 0.0011; 0s delay, p = 0.0553). Females did not reduce their responding until the 12s delay (p = 0.021) and continued to do so until the last delay (4s delay, p < 0.001; 0s delay, p = 0.0018). These results indicate temporal uncertainty induced by a schedule shift immediately caused differences in the degree of anchoring between schedules. Preferences for the large reward at all delays were higher when they were “anchored” by an initial 0 second delay than an initial 28 second delay. However, animals did show significant discounting on both schedules, as measured by changes in their preferences for the large reward (**Figure 2A**, main effect of delay, F_(4, 60.46)_ = 52.37, p < 0.001).

**Figure 2.**
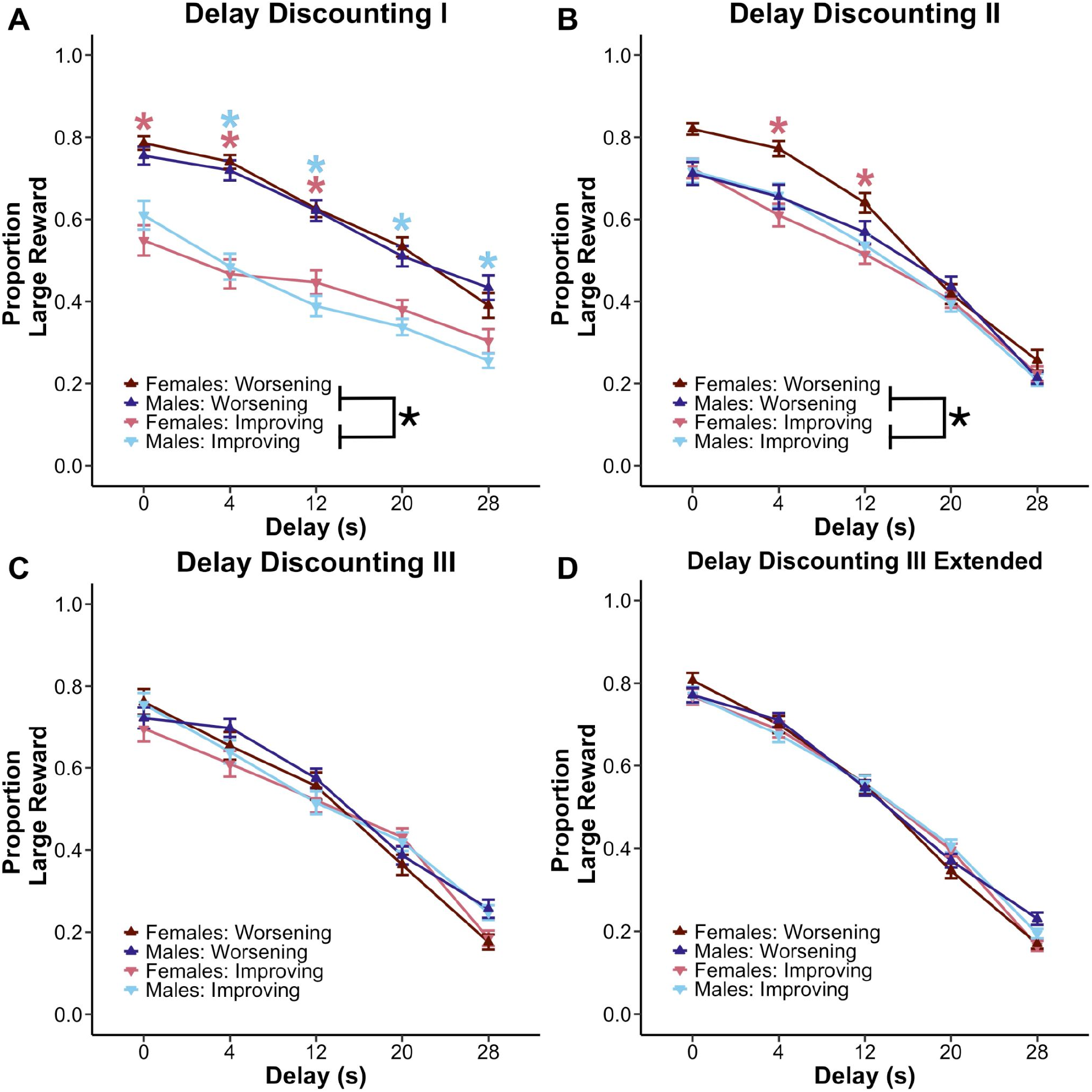
Anchoring effects in delay discounting are eliminated with extended experience. **A:** Mice of both sexes responded less for the large reward at the longest delays compared to the shortest, even with minimal experience on the tasks. However, there was a significant anchoring effect, such that mice in the Improving (started at 28s delay) condition had a persistent reduction in choosing the large reward compared to the Worsening (started at 0s delay) condition. **B:** Anchoring effects lingered into the second session of delay discounting, but were most pronounced in females. **C & D:** With continued experience, females and males reached similar discounting rates and no longer anchored their preference according to whether sessions started with a long or short delay. Figures depict mean ± SEM, black asterisks indicate significant anchoring effects (main effect of schedule) while colored asterisks indicate significant sex effect (planned posthoc comparisons of worsening and improving schedules within a sex at each timepoint) of p < 0.05.

Despite a second round of testing (**Figure 2B**, Delay Discounting II), anchoring effects persisted (**Figure 2B**, main effect of schedule, F_(1, 15)_ = 6.81, p = 0.0197). Again, sustained anchoring was suggested to be differently impacted in male and females (**Figure 2B**, sex x schedule x delay interaction, F_(4, 1251.92)_ = 5.21, p < 0.001). Female mice showed reduced responding on the Improving schedule at later delays (12s delay, p = 0.0081; 4s delay, p < 0.001), but not at the 0s delay (p = 0.0601). Female mice displayed a persistent anchoring effect compared to male mice. These data suggest that female mice may have increased sensitivity to anchoring effects, and/or increased sensitivity to unexpected changes in the task rules. Animals continued to show strong discounting to each transition of the delay (**Figure 2B**, main effect of delay, F_(4, 60.21)_ = 116.42, p < 0.001).

By the time animals were tested on Delay Discounting III (**Figure 2C**), there were no longer any anchoring effects or differences apparent in choice (p > 0.05). To establish that these preferences were stable, we continued testing the animals on each schedule for 9 days (performance across all days of Delay Discounting III in **Figure 2D**). Animals showed robust discounting to each delay (**Figure 2D**, main effect of delay, F_(4, 60)_ = 175.55, p < 0.001) that did not significantly differ according to whether they started the task with the longest or shortest delay, indicating stable discounting preferences. Taken together, our results suggest mice developed a steeper discounting function as they learned to anticipate the possible changes to the schedule. This change in anchoring is particularly evident in Improving schedules over time, suggesting an increased amount of sampling of the large reward as they learned that a long delay could improve. However, male and female mice seemed to make different types of adaptations.

### 3.2 Probability Discounting

#### Anchoring effects are persistent when discounting risky rewards

We put mice through probability discounting with “Worsening” and “Improving” schedules in order to challenge anchoring in response to uncertain large rewards. We tested mice on these schedules to see if risky rewards produced anchoring effects in a similar manner. If males and females are differently affected by uncertainty, it would stand to reason that those differences may be most reflected in the anchoring effects. We present the data from each round of Worsening and Improving schedules grouped together.

Unlike delay discounting, the first time mice experienced both the Worsening and Improving probability discounting schedules (Probability Discounting I), their preferences for the large reward across the entire sessions were not heavily anchored overall by the initial probability experienced (**Figure 3A**, no main effect of schedule, F_(1, 15.01)_ = 0.42, p = 0.527). However, mice showed schedule specific preferences depending on the probability (**Figure 3A**, schedule x probability interaction, F_(4, 1122.52)_ = 21.15, p < 0.001) and differences in the magnitude of anchoring between schedules (**Figure 3A**, sex x schedule interaction, F_(1, 15.01)_ = 4.83, p = 0.0441). Males specifically showed significant anchoring when the chance of winning risky rewards was at 25% (p = 0.0024) and 50% (p = 0.0144). Females however showed an increase in responding when risky rewards were guaranteed (100% chance, p = 0.0169). Male mice appeared to adjust choices more to losses than to wins compared to female mice. Regardless of sex, all mice showed significant discounting on both schedules, as measured by changes in their preferences for the large reward (**Figure 3A**, main effect of probability, F_(4, 60.88)_ = 123.43, p < 0.001).

**Figure 3.**
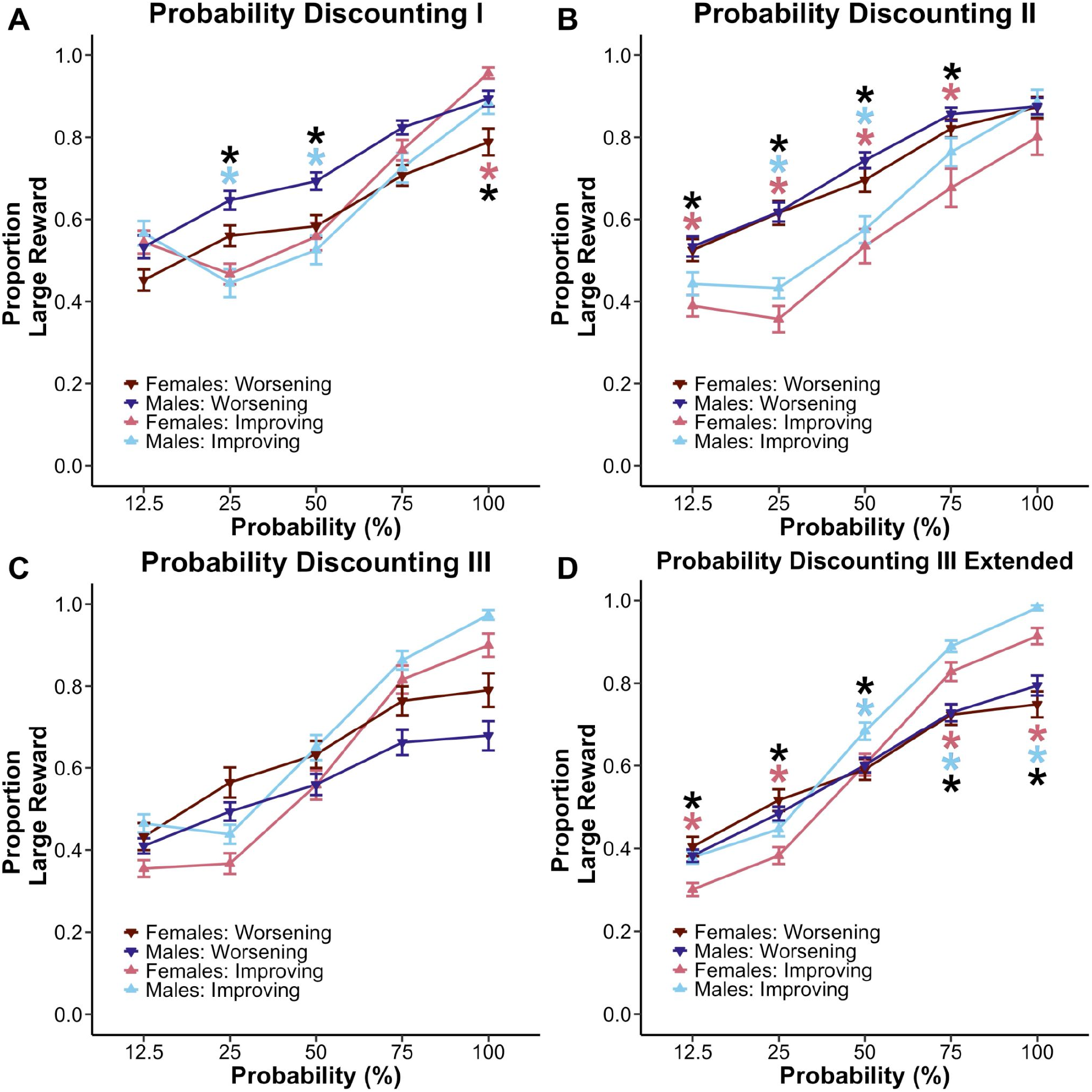
Anchoring effects continue to influence probability discounting behavior after extended experience. **A:** Regardless of uncertainty orientation, males and females choose the large reward less often with increasing risk. Interaction effects of schedule x probability were specific to the 25% (p < 0.001), 50% (p = 0.0195), and 100% (p = 0.0320) probabilities. **B:** For the second round of probability discounting, mice showed significant discounting of rewards and evidence of anchoring. Interaction effects of schedule x probability were specific to the 12.5% (p < 0.001), 25% (p < 0.001), 50% (p < 0.001), and 75% (p < 0.001) probabilities. Females were more sensitive to uncertainty where they showed anchoring effects starting at the 12.5% probability compared to males at 25% probability. **C & D:** With extended training, mice were sensitive to schedule effects at all probabilities: 12.5% (p = 0.0230), 25% (p < 0.001), 50% (p = 0.0410), 75% (p < 0.001), and 100% (p < 0.001) probabilities. Females showed decreased risky choice preference on the Improving schedule at risky probabilities. Male mice made specific adjustments only at safer probabilities. Figures depict mean ± SEM, black asterisks indicate significant anchoring effects (schedule x probability interactions) while colored asterisks indicate significant differences within a sex in responding to specific probabilities (planned posthoc comparisons) of p < 0.05.

As mice continued to gain more experience with the task design during the second round of testing (**Figure 3B**, Probability Discounting II), anchoring effects became more apparent (**Figure 3B**, main effect of schedule, F_(1, 15)_ = 37.74, p < 0.001). These anchoring effects again seemed to be specific at different probabilities of risky rewards (**Figure 3B**, schedule x probability interaction, F_(4, 1140)_ = 9.08, p < 0.001). Female mice significantly reduced their responding for risky rewards on the Improving schedule at 12.5% chance of reward (p = 0.0149) to the last probability (25% chance, p < 0.001; 50% chance, p = 0.0025; 75% chance, p = 0.0082). Male mice started to show reduced responding at 25% chance of reward (p < 0.001) and 50% chance of reward (p < 0.001). All mice continued to show strong discounting to each transition of probability (**Figure 3B**, main effect of probability, F_(4, 60)_ = 128.56, p < 0.001).

By the time animals were tested on Probability Discounting III (**Figure 3C/3D**), anchoring effects were still apparent (F_(1, 14.22)_ = 7.80, p = 0.0142). Mice again showed schedule specific preferences depending on the probability (**Figure 3D**, schedule x probability interaction, F_(4, 2198.20)_ = 45.12, p < 0.001) and differences in schedule preference (**Figure 3D**, sex x schedule interaction, F_(1, 14.22)_ = 4.98, p = 0.0422). Female mice reduced risky choices when uncertainty was high (12.5% chance, p = 0.141; 25% chance, p < 0.001) and increased risky choices when uncertainty was low (75% chance, p = 0.0136; 100% chance, p < 0.001). Male mice also made risk-based adjustments but only to maximize rewards when uncertainty was at or above chance level (50% chance, p = 0.0442; 75% chance, p < 0.001; 100% chance, p < 0.001). All mice showed significant discounting (**Figure 3D**, main effect of probability, F_(4, 60.57)_ = 315.46, p < 0.001). These results demonstrate that females and males made schedule specific adjustments in avoiding losses around an immediately risky schedule (i.e. Improving probability discounting) and that mice continue to remain sensitive to schedule effects with extended training.

### 3.3 Win-stay/lose-shift Adaptations to Risk Order

#### Differences in win-stay/shift behavior are reduced with extensive experience

Win-stay/lose-shift behaviors are important indicators of strategy specific adaptations to wins and losses. Win-stay ratios were calculated by dividing how often mice stayed on the same risky side after being rewarded divided by all rewarded risky responses. Lose-shift ratios were calculated by dividing how often mice switched to the small guaranteed side after not receiving a large risky reward divided by all losses on the risky side. We wanted to study if females and males made specific adaptations to wins and losses in response to different risk orientations, which can be a source of choice differences (Stopper et al., 2014).

At the beginning of probability discounting, mice seemed to win-stayed more on the Improving schedule (**Figure 4A**, main effect of schedule, F_(1, 15.043)_ = 15.03, p = 0.0015), but a sex x schedule interaction (**Figure 4A**, F_(1, 15.043)_ = 9.52, p = 0.0015) indicates only females win-stay more on the Improving schedule (p = 0.0075). Mice also lose-shifted more on the Improving schedule (**Figure 4B**, main effect of schedule, F_(1, 15.004)_ = 29.30, p < 0.001). Both male (p = 0.0071) and female (p = 0.0175) mice lose-shifted more in the Improving condition. These results suggest both male and female mice adjust their behavior to increased initial uncertainty in a similar way, but females specifically showed an increased ability to maximize rewards even when enduring great uncertainty.

**Figure 4.**
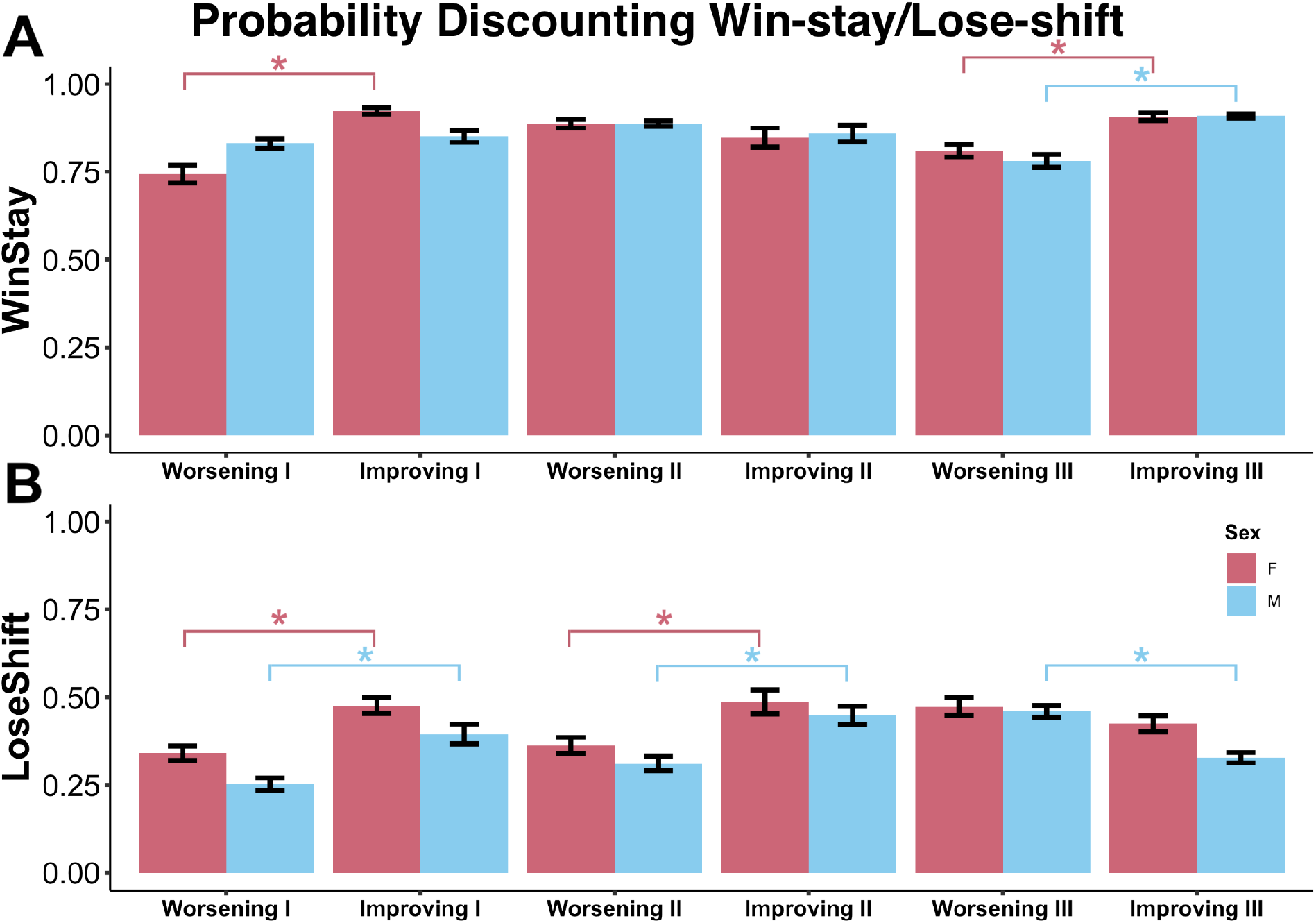
Adaptations to wins and losses with experience. We examined whether tendencies to win-stay/lose-shift explained sex differences in probabilistic responding. Win-stay ratios were calculated by dividing how often mice stayed on the same risky side after being rewarded divided by all rewarded risky responses. Lose-shift ratios were calculated by dividing how often mice switched to the small guaranteed side after not receiving a large risky reward divided by all losses on the risky side. **A:** Female mice specifically learned to increase win-stay behavior after their first experience with risky rewards. By the end of probability discounting, males and females learned to win-stay more on the Improving schedule. **B:** Male and female mice adapted to risky rewards by increasing their lose-shift behavior. By the end of probability discounting, only males showed lose-shift specific adaptations. Figures depict mean ± SEM, colored asterisks indicate significant within sex schedule effects (planned post hoc comparisons) of p < 0.05.

Mice made similar win-stay adaptations throughout the second round of probability discounting (**Figure 4A**), but continued to make loss specific adaptations (**Figure 4B**, main effect of schedule, F_(1, 255)_ = 32.15, p < 0.001). Increased lose-shift behavior on the Improving schedule was found again in both males (p = 0.0038) and females (p = 0.0128).

As mice finished probability discounting, male (p < 0.001) and female (p = 0.0069) mice continued to win-stay more on the Improving schedule (**Figure 4A**, main effect of schedule, F_(1, 14.967)_ = 48.434, p < 0.001). A main effect of schedule was found for lose-shift behavior (**Figure 4B**, F_(1, 14.493)_ = 19.026, p < 0.001), but only male mice exhibited decreased aversion to losses on the Improving schedule (p = 0.0024). Win-stay/lose-shift analysis revealed mice make specific schedule adaptations to wins and losses and those adaptations remain pervasive until the end of discounting.

### 3.4 Computational models of choice variability in Delay and Probability Discounting

#### Overall decision noise decreases in probability discounting compared to delay discounting

Win-stay/lose-shift analyses revealed some learning specific effects, but did not explain changes in preference over renditions of the task. We posited that snapshot win-stay/lose-shift analyses might not capture broader trends driving choice behavior. There is a growing amount of evidence suggesting exploration is a key latent variable driving choice behavior (Ebitz et al., 2018; Ebitz et al., 2019). Therefore, we pursued two discounting models for delay and probability, each incorporating a value parameter (*k* for delay, *h* for probability; Green et al., 2014) and an inverse temperature parameter capturing variability in choice around these value preferences (β). This allowed us to track adaptations in value and choice within and between tasks. We excluded sex as a factor to increase power and better detect schedule specific adaptations.

Analysis of the delay discounting rate parameter revealed mice had smaller discounting rates for the Worsening condition (**Figure 5A**, main effect of schedule, F_(5, 75)_ = 8.039, p < 0.001). Mice had a steeper discounting rate initially on the Improving condition compared to the Worsening condition (**Figure 5A**, p < 0.001), possibly due to the introduction of uncertainty induced by a switch in orientation. Probability discounting rates were also schedule dependent (**Figure 5A**, F_(5, 75)_ = 7.706, p < 0.001), especially for the second session of discounting (**Figure 5A**, p = 0.0086). However, these value parameters did stabilize for the last rounds of delay and probability discounting.

**Figure 5.**
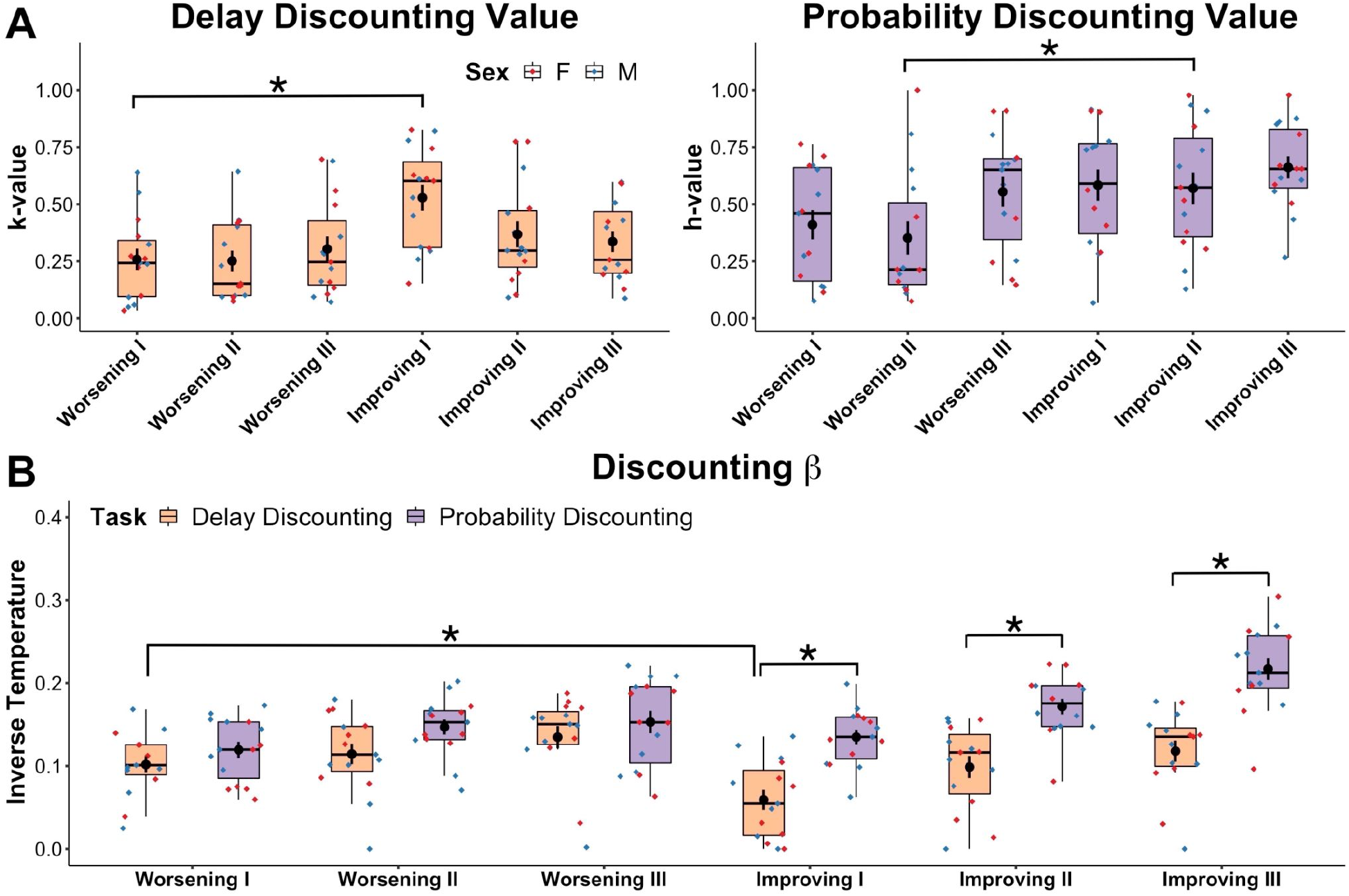
Delay promotes flexibility, probability promotes rigidity. We modeled discounting rates (*k* and *h*) and decision noise (inverse temperature, β) for each mouse across all days of testing. **A:** Delay discounting *k*-values and probability discounting *h*-values show relatively few changes in discounting value estimates over these schedules, especially past initial experiences, despite large changes in actual discounting behavior. **B:** Inverse temperature (β, reflecting decision noise) parameters for delay and probability discounting reveal large changes in the noisiness of choices as schedules change. Inverse temperature increased over time with extended experience with both tasks, indicating decreased decision noise in choices with experience. Risky rewards produced by probability discounting produced the largest beta value, indicating the closest adherence to choosing the most valued option. Figures depict individual points for males (blue) and females (red) and box-and-whisker plots with mean ± SEM overlaid, asterisks indicate significance of p < 0.05.

Between-task analyses of the inverse temperature in our discounting model revealed a general delay versus probability effect where mice tended to explore options more when rewards were uncertain (**Figure 5B**, main effect of task, F_(1, 15)_ = 25.096, p < 0.001). Mice increased repetitive choices in probability discounting compared to delay discounting when comparing session one (**Figure 5B**, F_(1, 15.021)_ = 28.526, p < 0.001), session two (F_(1, 15)_ = 21.914, p < 0.001) and session three (F_(1, 15)_ = 12.775, p = 0.0028). Further investigation into these effects revealed that mice repeated choices significantly more when initial uncertainty was high compared to when initial delay was high. Mice engaged in increased repetitive choice on Improving I (**Figure 5B**, p < 0.001), Improving II (p = 0.001), and Improving III (p = 0.001).

Within-task comparisons of the choice parameter revealed mice adapt to initial temporal uncertainty caused by an orientation switch through increased sampling of reward options (**Figure 5B**, Improving I < Worsening I, p = 0.0106). With extended training, mice primarily mitigate the use of repetitive choices on probability discounting. Between and within subject analyses overall show a dissociation of the processing of delayed and probabilistic rewards where mice adjust to high temporal uncertainty by increasing decision noise, but decrease decision noise when probabilistic uncertainty is high (i.e. the Improving conditions).

## 4. Discussion

We trained male and female mice in a novel battery of delay and probability discounting schedules, in order to assess 1) if mice exhibit stable choice behavior across these tasks, and 2) if sex differences affected stabilization of choice behavior. These tasks are high-priority goals for cross-species translation, and there is some controversy over whether these two tasks test similar or distinct constructs. Overall, we found mice showed substantial discounting, indicating sensitivity to the structure of both tasks. Mice formed stable choice behavior in delay discounting over time, but continued to show anchoring effects throughout probability discounting. To understand why there might be differences in the persistence of anchoring effects between these tasks, we examined win-stay/lose-shift strategies in probability discounting, and across both tasks, fit discounting models to animal data that included both value parameters (*k*/*h*) and an inverse temperature parameter to capture decision noise (β). Win-stay/lose-shift analysis hinted at the presence of schedule-dependent shifts in choice, similar to the anchoring effects prominent throughout probability discounting. Discounting models revealed that mice learned throughout training on both tasks to reduce decision noise, leading to demonstrations of stabilized reward preference, without changes in the discounting parameters themselves. Our results indicate exploratory decision noise may be underappreciated contributors to behavior in animal models in reward-guided decision making tasks.

Human discounting analyses use value-based models (i.e. *k* and *h* values) to determine the extent of discounting behavior according to the value of the reward. Discounting research has demonstrated how hyperbolic models of value best explain discounting behavior of both humans and animals (Green et al., 2014; Vanderveldt et al., 2016). Discounting steepness is believed to follow a value rule that is liable to change in response to individual factors such as risk tolerance and choice impulsivity (Odum, 2011b; Simon et al., 2009). Our results are in line with previous research where modelling around a value parameter describes stable discounting behavior (Green et al., 2014; Odum, 2011a). However, recent research into animal discounting shows that animals do not always optimize for the discounted value of the reward and implies discounting can arise from multiple individual sources (Blanchard et al., 2013; Hayden & Niv, 2020). In our data, adding an inverse temperature (β) parameter substantially improved our model fit. In other words, it isn’t that mice appeared not to use a value rule (*k*/*h*), but that their adherence to a value rule when making decisions changed dramatically over training. Adherence to the value parameter was higher in delay discounting than in probability discounting, and we saw a greater persistence of anchoring influences in probability discounting even after many weeks of experience. Our modeling results combined with our win-stay/lose-shift analysis indicated that increased uncertainty experienced in probability discounting tasks may have promoted increased repetitive behavior, consistent with previous rodent literature (Derusso et al., 2010). The idea that task-based uncertainty can promote less flexible choice helps explain how decreased decision noise in probability discounting could reflect anchoring choices to reward probabilities experienced earlier in the schedule. Our choice to add a decision noise parameter allowed us to better capture adaptations to schedules across tasks and provide an explanation for differences in choice preference for both tasks. This suggests that decision noise is an underappreciated contributor to value-based decisions in animal models.

Sex effects have not been consistently observed in reward-guided delay or probability discounting tasks (Grissom & Reyes, 2019; Orsini & Setlow, 2017). Sex differences appear in discounting tasks depending on the type of uncertainty or consequences (especially aversive outcomes; (Orsini et al., 2016; Orsini & Setlow, 2017)), but generally do not appear when the risk is the loss of a reward (Grissom & Reyes, 2019). In the current dataset, animals were able to learn a consistent pattern of behavior to delay regardless of the order of presentation, but probability discounting did not lead to a consistent pattern of choices across sexes. Despite differences in the degree of anchoring across probabilities seen in males versus females in our choice data (Figure 3 and 4), we were unable to capture those effects in our computational model (Figure 5). It is possible that we were underpowered to detect sex differences in modeling decision noise given that our lab has observed it in the past (Chen, Knep, et al., 2020). Sex differences might also be better captured with an additional latent variable not defined within our model.

While delay and probability discounting are often both thought to measure an aspect of choice impulsivity (Dalley et al., 2011; Green & Myerson, 2004; MacKillop et al., 2016; Strickland & Johnson, 2020), they are found to be weakly correlated even in humans (Green et al., 2014; Strickland & Johnson, 2020). As such, straightforward value models may miss hidden traits specific to delay or risk. In attempting to model these tasks in rodents, one source of variability in reward preference could arise from differences in choice patterns outside of the optimal choice. We included a decision noise parameter in order to capture decision noise hidden in the value parameters (Nussenbaum & Hartley, 2019). Our results suggest mice not only make value-based decisions, but they also adapt decision noise around the discounted value of a reward. Differences in choices across schedules are better explained by changes in decision noise as opposed to the value parameter. Decision noise is therefore a significant contributor to discounting behavior in our mice, and may be an underrecognized contributor to rodent choice behavior in other contexts.

While we designed our tasks to capture adaptations in value and decision noise, some caveats come with the sequential design of our tasks. Delay and probability discounting rates are significantly impacted by the order of presentation of delayed or probabilistic uncertainties and could also be affected by whether animals experience delay or probability discounting first (Rung, Frye, et al., 2019; St Onge et al., 2010; Tanno et al., 2014). It is worth noting mice did stabilize in their performance in delay discounting, despite experiencing two different orders of delay. The order of exposure to delay and probability discounting could account for some of the persistent anchoring we observe with probability discounting. Our delay and probability discounting tasks are structurally similar and thus could influence choice behavior acquired through probability discounting (Neville et al., 2020). We did, however, use a novel house light cue introduced during probability discounting to help mice distinguish between both tasks. Further, each task promotes different types of uncertainty (Garr et al., 2020) which provides an opportunity for new learning. Our choice and computational data support these ideas as mice showed differences in adaptations across both tasks. Still, we can’t fully rule out order effects, but this seems less likely given that our behavior and modeling parameter fits indicate that animals continued to adjust their behavior across the duration of the probability discounting task.

Our results support a role for computational modelling in identifying latent variables that contribute to decisions in rodent tasks. As noted above, while *k* and *h* parameters can be used to reflect value in discounting tasks, they are not able to capture the contributions of other variables that might influence choices. Probabilistic tasks, including discounting, are amenable to analyses of choice patterns following wins and losses (e.g. win-stay and lose-shift) but lack a parallel in delay discounting. A global parameter to compare decision noise across tasks is important when assessing choice behavior and testing for factors contributing toward impulsive behavior (Dalley et al., 2011; MacKillop et al., 2016; Strickland & Johnson, 2020). The inverse temperature parameter we included in our computational model helps bridge the gap between the two tasks by allowing for task comparisons and examining influences of order presentation and group differences in valuation of rewards. Our results demonstrate how computational models account for decision noise are better to detect different sources of behavioral variability, such as anchoring effects, demonstrated in sequential versions of decision making tasks.

## Acknowledgements

We would like to thank Jake Jeong and Jack Leschisin for expert technical assistance with behavioral experiments. This work was funded by NIMH R01 MH123661, NIMH P50 MH119569, NIMH T32 training grant MH115886, and startup funds from the University of Minnesota.

## Author Contributions

GRR and ATH performed experiments and curated the data. GRR and CSC conducted analyses, validation of analyses, and data visualization. GRR wrote the original draft, and GRR, LSCP, CSC and NMG reviewed and edited the manuscript. NMG conceptualized the project, acquired funding, and supervised the project.

